# Performance Evaluation of Traditional Signal Processing Methods in Localizing *Tursiops truncatus* Whistles in a Reverberant Aquatic Environment

**DOI:** 10.1101/642736

**Authors:** SF Woodward, MO Magnasco

**Affiliations:** Laboratory of Integrative Neuroscience, Center for Studies in Physics and Biology, The Rockefeller University, New York, NY, United States

## Abstract

Relative to individually distinctive *signature whistles*, little is known about the “non-signature” calls – particularly the non-signature whistles – of the common Atlantic bottlenose dolphin, *Tursiops truncatus*. While such calls are suspected to serve social function, tracking their exchange among conspecifics and correlating their usage with non-acoustic behavior has proven challenging, given both their relative scarcity in the dolphin repertoire and their characteristic shared use among dolphins, which precludes the unique identification of callers on the basis of whistle properties alone. Towards the goal of robustly identifying the callers of non-signature whistles (equivalently, attributing non-signature whistles to callers), we present a new, long-term audiovisual monitoring system designed for and tested at the Dolphin Discovery exhibit of the National Aquarium in Baltimore, Maryland. In this paper, we confirm the system’s ability to spatially localize impulse-like sounds using traditional signal processing approaches that have already been used to localize dolphin echolocation clicks. We go on to provide the first rigorous experimental evaluation of the component time-difference-of-arrival-(TDOA) extraction methods on whistle-like tonal sounds in a (reverberant) aquatic environment, showing that they are generally not suited to sound localization. Nevertheless, we find that TDOA extraction under these circumstances is performed significantly better using a Generalized Cross-Correlation with Phase Transform (GCC-PHAT) method than a standard circular cross-correlation method, a potentially important result.

## Introduction

Ever since mid-20th century studies [1, 2] observed that Atlantic common bottlenose dolphins (*Tursiops truncatus*) use a variety of tonal and burst-pulse calls, researchers have explored the possibility that dolphins (and specifically the common bottlenose) manifest communication mediated by sound. While Dreher [2] provided early evidence that even a single dolphin can use multiple distinct tonal sounds, termed whistles, later Melba and David Caldwell [3, 4] famously observed that each common bottlenose dolphin shows significant preference for a single, individually distinctive whistle, which they termed its *signature whistle*. While it would become apparent that the strong signature whistle preference that the Caldwells observed applies primarily to individuals isolated from conspecifics (though a weaker preference is still observed for individuals in close proximity), signature whistles would nonetheless become the focus of many research efforts in exploring the communicative utility of whistles in *T. truncatus*; little would be learned about non-signature whistles by comparison [5].

Reasons that signature whistle studies continue to outnumber non-signature whistle studies include not only signature whistles' numerical prominence in dolphin recordings from both the wild and captivity, and particularly from convenient capture-and-release studies [6], but also the relative ease with which specific instantiations of signature whistles can be paired with originating dolphins, even in social settings. The use of a particular signature whistle implies the identity of the caller [5] (while signature whistle copying does occur, it seems to occur only with modification of the original whistle [7]). This association between a call and its caller, which we term sound or whistle attribution, allows for the correlation of a call with gestural and other non-acoustic behavior [5, 6], and potentially for the examination of acoustic exchanges among conspecifics [5, 8]. In contrast, non-signature whistles are relatively scarce, and by definition are not individually distinctive, which precludes such trivial whistle attribution. Without achieving robust non-signature whistle attribution, it will remain impossible to study non-signature whistles to the degree signature whistles are, or to gain a comprehensive understanding of bottlenose dolphin communication.

This paper is broadly concerned with the challenge of performing whistle attribution within an audio recording of multiple socializing dolphins, which is relevant to the functional study of not only non-signature whistles, but potentially of non-whistle calls (e.g., burst-pulse sounds). Highlighting the significance of this challenge, it is notable that no studies exist claiming to attribute whistles in a group of more than two dolphins reliably enough to accommodate a sequence analysis of non-signature calls. More specific than the the challenge of whistle attribution, this paper is concerned with the difficulty of performing whistle localization, the task of determining the physical coordinates of the origin of a whistle. Explicitly or implicitly this is often a prerequisite of sound attribution, preceding a step that matches the obtained coordinates to visual (e.g., video) identities.

With exceptions, to be mentioned, sound localization often involves obtaining time-differences-of-arrival (TDOA’s) for a sound (or signal) of interest between several pairs of sensors, and solving the corresponding nonlinear, non-convex system of geometric equations using one of several approaches [9] – in this paper we use a reliable solution termed Spherical Interpolation, which is an optimal estimator under the assumption of Gaussian error [9–12].

Intuitively, the best method for obtaining TDOA’s can depend heavily on the nature of the signal of interest. TDOA’s for strongly-peaked, pulse-like sounds can often be obtained by simply thresholding the signal amplitudes to find signal onset times, with TDOA’s then obtained by subtraction; echolocation click TDOA’s have been successfully obtained in this way [13–15]. TDOA’s for signals that are relatively extended and heterogeneous in time, which describe whistles, are often obtained by cross-correlation-based approaches [16–18] that rely on finding the time delay that corresponds to the optimal overlap between received signals of two different sensors. To find the TDOA, *t*_*delay*_, for signals *r*_*i*_(*t*) and *r*_*j*_(*t*) from sensors *i* and *j*, respectively, the simplest cross-correlation-based approach searches for a unique sharp peak:

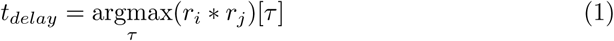

Working to localize bottlenose whistles, other authors have at best achieved modest results in irregular, low-reverberation environments [18–23]. While a review of the relationship of all methods to the above framework would be out of the scope of this paper, we note that many of the previous authors and others [16] have noted difficulty localizing dolphin whistles in reverberant environments. As has been analytically described [16], the standard cross-correlation-based approaches are potentially disrupted in environments where the original signal becomes stacked with copies of itself resulting from reflections, or generally *multipath effects*. In this case, no single peak in the cross-correlation will exist, and the largest peak will not necessarily correspond to the desired pair of direct sound paths. No modification of the cross-correlation has been proven robust to multipath effects, however one established modification of the cross-correlation that has been suggested to be resilient is the Generalized Cross Correlation with Phase Transform (GCC-PHAT) [17], which we evaluate here.

We note that promising new approaches to whistle attribution exist that are not directly addressed in this paper. These include a sound-localization-based method termed SRP-PHAT (a beam-forming, cross-correlation-based method), which has achieved modest success (40% recall) attributing whistles in a low-reverberation environment [18]; tag-based methods of sound attribution [24–26]; and machine-learning-based approaches, which we have proposed elsewhere [27]. The first method, relying on cross-correlation, is theoretically susceptible to the same complications introduced by the multi-path problem discussed earlier, and has not been tested in a highly reverberant environment.

The goal of this paper is to provide an experimental performance evaluation of the more common forms of sound localization on whistle-like sounds. The first part of this paper is concerned with showing that our custom system of 16 permanent hydrophones, located at the Dolphin Discovery exhibit of the National Aquarium in Baltimore, Maryland, is capable of localizing ideal, pulse-like sounds. The second part of this paper is concerned with attempting to employ similar methods to attempt localization of whistle-like sounds, also originating from known locations in the pool.

## Materials and methods

### Overview

We obtained acoustic and visual data from equipment deployed at the Dolphin Discovery exhibit of the National Aquarium in Baltimore, Maryland. The exhibit’s 110’-diameter cylindrical pool is subdivided into one approximate half cylinder, the *exhibit pool* or EP, as well three smaller holding pools, by thick concrete walls and 6’x4.25’ perforated wooden gates; all sub-pools are acoustically linked. The data were obtained from the EP, when the seven resident dolphins were in the rear pools.

The basic experimental setup for obtaining acoustic data involved both input and output subsystems, which shared two synchronized, poolside MOTU 8M audio interfaces connected by a fiber optic cable to a Mac Pro in the dolphin amphitheater sound booth.

The output subsystem transmitted Matlab-generated sounds through the MOTU interfaces to an omnidirectional marine Lubbell LL916H speaker. The speaker was secured at known heights below a modified marine-buoy-based flotation device, which could be moved across the surface of the EP using four ropes, which were secured to the flotation device as well as four poolside attachment points. An optical target mounted to the buoy allowed the surface coordinates of the buoy to be determined using four Bosch 225 ft. Laser Measure devices and a straightforward triangulation procedure.

All output sounds were played at 14 locations inside the pool. The 14 locations corresponded to 7 unique positions on the water surface on a 3 × 5 cross, at 6 feet and 18 feet deep. Approximate surface positions are shown in Fig 1; the difference between adjacent horizontal and vertical positions was 10-15 feet. The speaker could sway from its center point by as much as a few feet during calibration.

**Fig 1.**
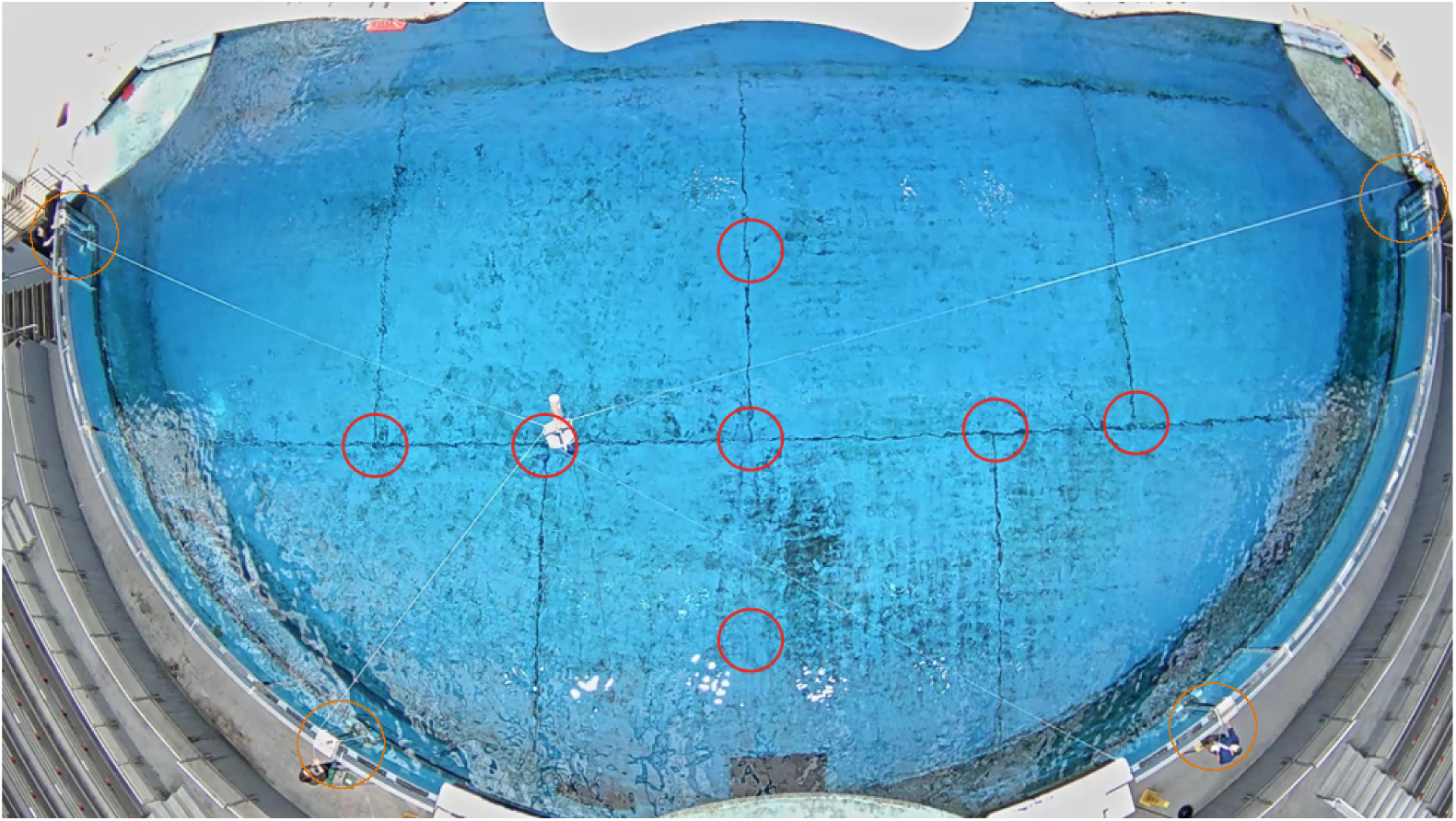
Pool test point and hydrophone array layout. The National Aquarium Exhibit Pool (EP) is shown, as visualized by the overhead AXIS P1435-LE camera. Circled in red are the approximate surface projections of the fourteen points at which sounds were played (i.e., seven distinct points). Circled in yellow are the four hydrophone arrays, each containing four hydrophones.

Also at the four poolside attachment points for the ropes were underwater sound receivers. Specifically, at each of the four locations was a large custom “hydrophone array,” designed to suspend four standard hydrophones (SQ-26-08’s from Cetacean Research Technology, with approximately flat frequency responses between 200 and 25,000 Hz) approximately six feet below the water surface in such a way that they were effectively isolated from the resident dolphins. While removable, the arrays (Fig 2) underwent an exhaustive engineering and testing process, and proven robust to long-term exposure with minimal maintenance. Through a series of connections, all sounds were ultimately recorded to the Mac Pro cited above.

**Fig 2.**
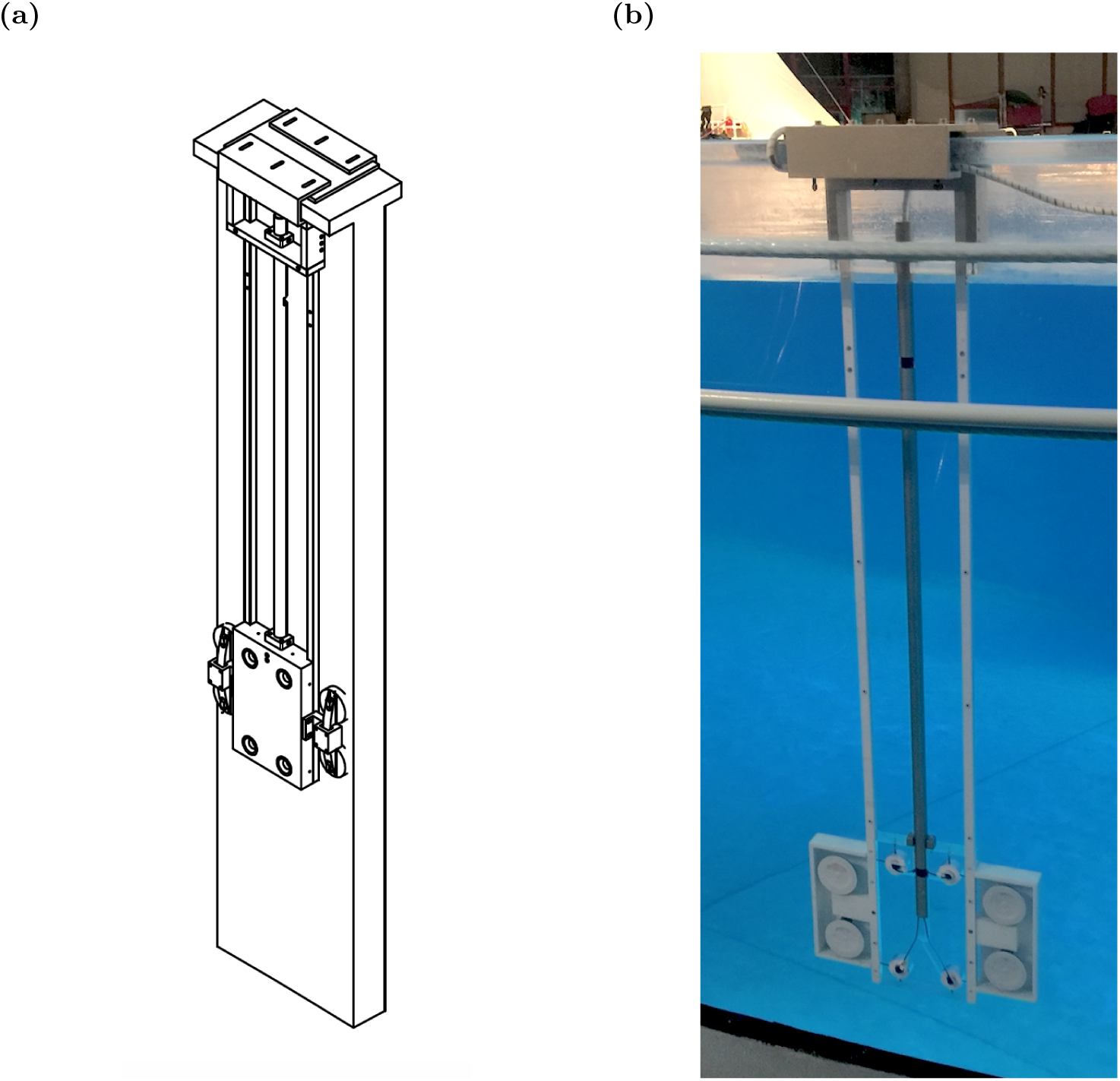
Hydrophone Array. (a) Selection from a mechanical drawing. Array is depicted attached to a small section of acrylic wall, which visibly has an approximate “T” cross-section. (b) A photo of an installed array.

The study also made use of a central AXIS P1435-LE camera, managed by Xeoma surveillance software and custom Matlab code from the same computer.

### “Snow White” Noise, Impulse Response Function Localization

One of the fundamental tools used to understand and account for a space’s acoustic properties is the impulse response function (IRF) [28]. Not only can the impulse response be analyzed to obtain information about a space’s acoustic properties between a specific source and receiver pair (such as values describing the nature of reflection/reverberation, and potentially clues to where boundaries are located), but, for a linear, time-invariant system, the IRF provides a complete description of an arbitrary signal’s transformation between the source and receiver. For such a system, the received signal *y*[*t*] for a known source signal *x*[*t*] can be described as a convolution with the IRF, *h*[*t*]:

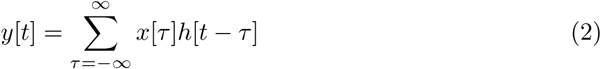

We understand that this relation is unique for every unique source-sensor pair (especially with regards to spatial positioning). This relation simplifies in Fourier space, and can be rearranged to solve for the IRF:

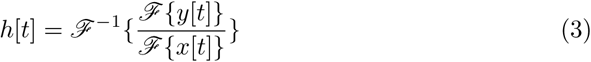

The above suggests that we can obtain the linear impulse response given any pair of source and received signals. However, to ensure that the denominator is nowhere zero, and to avoid biasing for any frequency in particular, in practice it is best if the source signal’s power is uniformly distributed across all frequencies (also a property of a true impulse). Such a signal can be obtained by inverse Fourier transforming a signal designed in complex frequency space that has unitary power at all frequencies, with random phase. This is the basis of what we term the *snow white* method of sound calibration. Note that if the signal thus obtained is not played at the appropriate sampling rate in its entirety, its power spectrum will not be unitary, but rather random with powers falling on a Gamma distribution – this is consistent with the power reflecting the absolute value of Gaussian variable pairs in real and imaginary space. The duration of the signal should be longer than the longest expected multipath travel time (one second was used). Moreover, the signal can be repeated a number of times (360 repeats, equivalent to 6 minutes was used) to account for various stochastic effects: the IRF is constructed from the median value for every time point. The sound was played at each of 14 locations, the received signal divided by the the source signal as shown in Equation 3 to obtain an IRF for each of the 16 × 14 sensor-location pairs. We settled on an appropriate amplitude by considering the strength of test data along with husbandry concerns.

The primary use of the IRF’s thus obtained was to determine how well a standard method of sound source localization, Spherical Interpolation [9–12], can localize a real, low-noise impulse in the National Aquarium EP; this would represent a best case scenario for how well other sounds can be localized using this method. Moreover, this analysis might provide a reasonable approximation of how well impulse-like common bottlenose clicks can be localized with this method using our particular experimental setup.

For *N* sensors, Spherical Interpolation requires *N* − 1 time-differences-of-arrival (or TDOA’s); these *N* − 1 TDOA’s correspond to the arrival time differences between *N -* 1 unique hydrophones and a single designated reference hydrophone. In our case, for 16 hydrophones, 15 arrays of TDOA’s were generated for a given impulse, one array for each of 15 hydrophones paired with one common reference; each array was of length 20,000. Letting integers *k* ∈ {1, …, 20000} index array position, and integers *i* and *j* index hydrophone and pool test location, respectively, each value *z*_*ijk*_ was drawn from

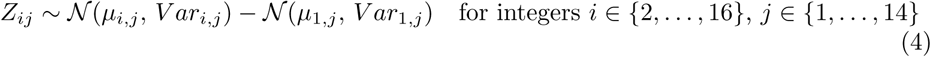

where *µ*_*i,j*_ indicates the time corresponding to the peak amplitude of the IRF’s first incidence in hydrophone *i* at location *j*, and *V ar*_*i,j*_ approximates the variance in these times among hydrophones in the same panel (which should be effectively the same at our temporal resolution). The (15 × 20,000) matrices of TDOA’s, one for each pool test location, were used to compute source location point clouds. These clouds were approximately ellipsoid, and their major radii, minor radii, areas, and center displacements from the true testing location were measured.

Separately, the absolute deviation of measured arrival times of the IRF’s from theoretical arrival times were used to visualize the relative error across hydrophones.

### Whistle-Like Sounds, TDOA Extraction

As described previously [27], we also played 128 unique sounds (analysis performed on 127) whose parameter values approximate those of *T. truncatus* whistles. These sounds were generated with amplitudes matched to the sound amplitudes of whistles used by the pool’s resident dolphins. In total, 1,605 recorded tones were successfully extracted for analysis.

The initial goal was to localize the tonal, dolphin-whistle-like sounds in the same way as IRF’s. However, we would discover TDOA’s suitable for localization were not obtained using the extraction methods considered. Thus, our analysis of the localization pipeline for tonal sounds focused on the accuracy with which TDOA’s could be obtained.

We used a variety of methods to obtain TDOA’s (or the component single-hydrophone arrival times). These methods included a custom thresholding algorithm for consistently locating whistle arrival times based on waveform features, locating TDOA’s by identifying the maxima of circular cross-correlation between pairs of hydrophone waveforms (both a custom implementation and the Matlab “finddelay” function were tested), locating TDOA’s by identifying the maxima of generalized cross-correlation with phase transform (GCC-PHAT) [17, 29], and locating TDOA’s by identifying maxima of a two-dimensional cross-correlation of spectrograms.

When employing the various cross-correlation-based methods of obtaining TDOA’s, we used both a single hydrophone signal and the original source signal; the latter operation results in so-called matched filtering. Moreover, prior to computing TDOA’s we applied either a “tight” or “loose” bandpass filter to the signals; the boundaries of the former were offset by 250 Hertz of the original signal, and boundaries of the latter were offset by 1000 Hertz.

The TDOA’s found using the various techniques were compared to theoretical values, calculated from known play locations. We considered the mean of the deviations (raw values), the mean of the absolute-value deviations, and the mean of the absolute-value deviations with 20% outlier truncation along the hydrophone axis, in case a method happened to fail on a small subset of received signals.

In an attempt to de-noise the received tonal signals of multipath effects, yielding signals more amenable to TDOA extraction with the removal of false peaks from the cross-correlation (see Introduction, Equation 1), we also de-convolved the tonal signals with the IRF’s previously obtained. We implemented the Wiener deconvolution, which in certain circumstances reduces the noise observed with the standard deconvolution. Our analysis was limited to a qualitative, visual examination of the signals, of which we present a sample.

## Results

### IRF Localization

Fig 4 shows the results of IRF localization in seven plots, one for each location (only the upper seven testing locations are shown; the bottom are similar). While these plots are two-dimensional projections of three-dimensional results, significant information is not lost in the projections, as the hydrophone panels have effectively no localization precision along the Z-axis – an expected result of Z-axis hydrophone placement limitations. In two dimensions the clouds of localized points can be approximated as ellipsoids and characterized by major and minor radii. Under this approximation, we have calculated the average areas of the clouds and the percentage of the EP they occupy, as well as the distance between the true calibration points from the nearest cloud points; this is done grouping all calibration locations as well as separately grouping midline and non-midline locations. These data are in Table 1. As is also visible from the plots, the midline group is localized more poorly, likely a consequence of the array and pool geometry that requires further examination.

**Table 1.** Performance of Spherical Interpolation on IRF Signals.

| Statistic | Value |
| --- | --- |
| Mean Cloud Area (ft<sup>2</sup>, % of pool) | 10.84 +/- 9.63 (0.24 +/- 0.21) |
| Mean Cloud Area, non-midline (ft<sup>2</sup>, % of pool) | 6.310 +/- 2.51 (0.14 +/- 0.054) |
| Mean Cloud Area, midline (ft<sup>2</sup>, % of pool) | 16.88 +/- 12.48 (0.37 +/- 0.27) |
| Mean Distance of True Point to Cloud (ft) | 2.63 +/- 3.76 |
| Mean Distance of True Point to Cloud, non-midline (ft) | 2.29 +/- 1.75 |
| Mean Distance of True Point to Cloud, midline (ft) | 3.08 +/- 5.86 |

**Fig 4.**
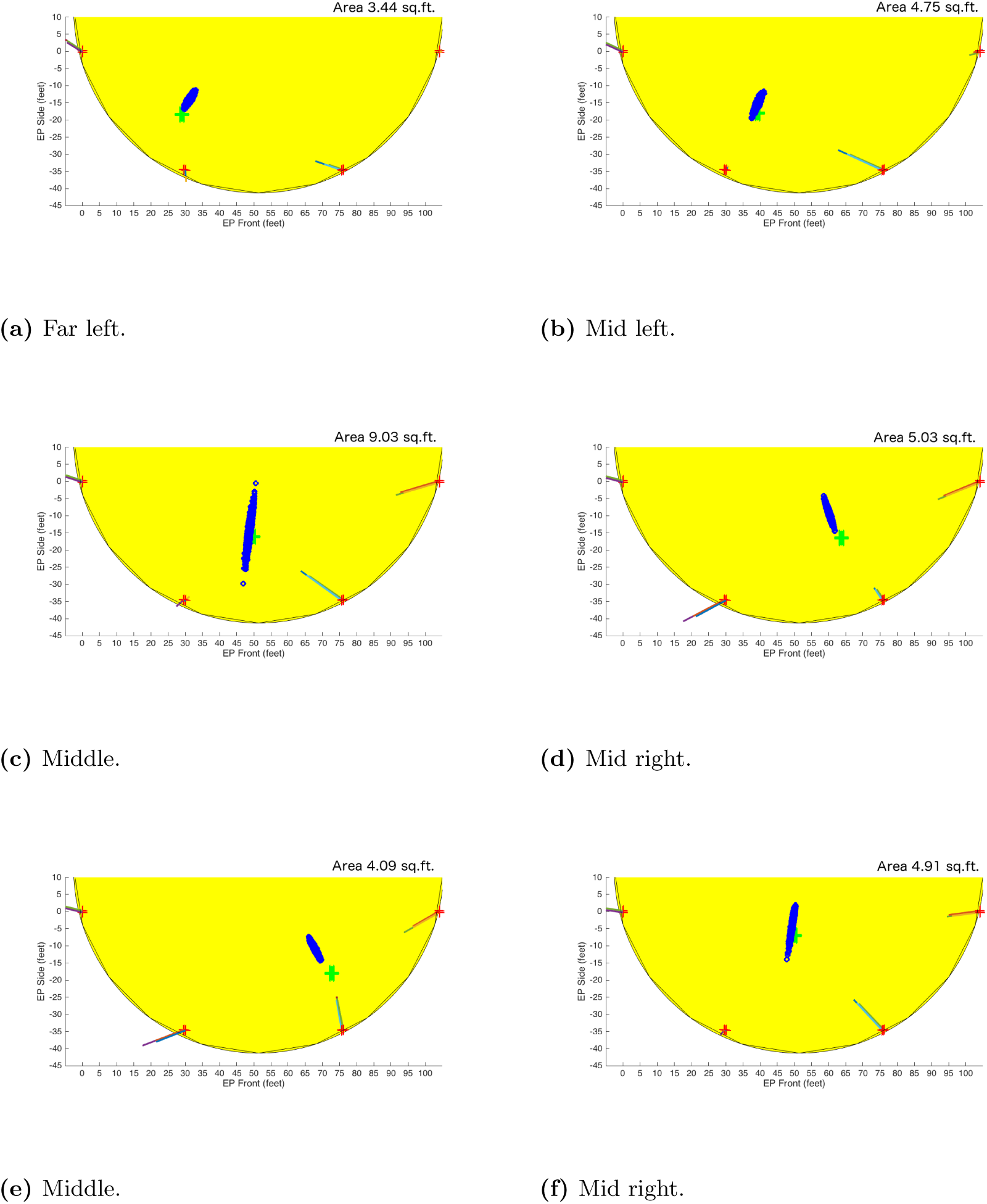
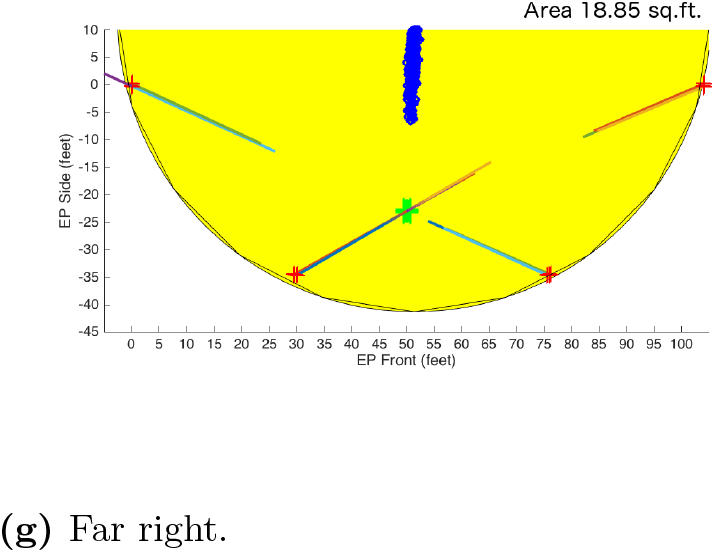
Each plot represents a simplified overhead 2D projection of the EP. The true position of the IRF is indicated by a green cross. Each blue asterisk represents the estimated position of the IRF based on estimated time of arrivals with Gaussian noise, described in the text. The red crosses around the EP perimeter indicate hydrophone positions. The lines drawn from them are proportional to the hydrophones’ estimated time estimation error; when oriented towards the true point, they indicate that estimated times were too late; when towards, too early. Plots are shown for 7 locations.

The data indicate that the cloud of localization points consistently occupies less than 1% of pool area (or, equivalently in this case, volume) – note that the plot markers are somewhat exaggerated in size for visibility – and that the true sound source is reliably within 5 feet of it. There is no appreciable overlap of clouds belonging to unique calibration locations in XY except at midline positions, where distinguishing among the calibration points is difficult.

The estimated time delays of the IRF’s were also used to determine whether any of the 16 hydrophones consistently underperforms. For each calibration point, the ideal arrival time differences were calculated (requiring a knowledge of the speed of sound and the location of source and hydrophones), and deviations from the estimated arrival time differences calculated. The mean and standard deviations across all calibration points were calculated for every hydrophone and are plotted in Figure 5. No significant difference among hydrophones is apparent.

**Fig 5.**
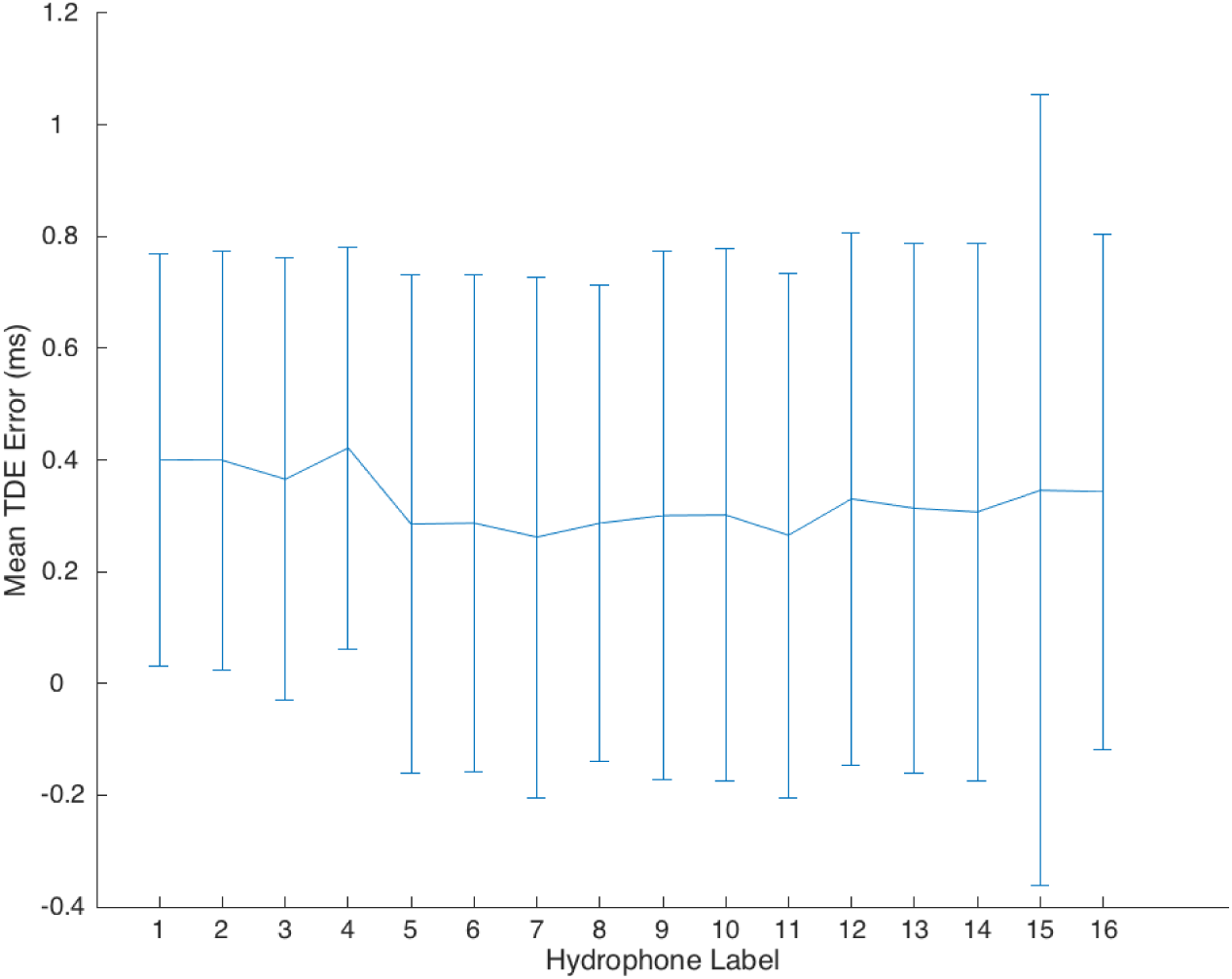
Mean Time Delay Estimation (TDE) Error of Hydrophones Plot of mean Time Delay Estimation (TDE) error of hydrophones, calculated from estimated and theoretical time delays for IRF’s. Error bars indicate standard deviations.

### Whistle-like Sound TDOA Extraction

Tables 2-5 show the performance of the various methods for TDOA extraction. As it is readily apparent from Fig 6 that the different groups differ greatly in variance, we favored nonparametric statistical tests as much as possible. We performed a Kruskal-Wallis test on the deviations of the true TDOA’s from the ideal TDOA’s, which indicated that the deviations were unlikely to be drawn from the same distribution (p = 0.0069); however, comparison of pairs of the corresponding mean rank intervals indicated that no two groups have statistically different means, in a territory around zero. A Kruskal-Wallis test performed on the absolute value of the deviations of the true TDOA’s from the ideal TDOA’s indicated that the absolute value deviations were unlikely to be drawn from the same distribution (p ≈ 0); more importantly, the post-hoc analysis indicated that the methods fell into distinct groups based on their mean rank intervals, shown in Fig 7.

**Table 2.** Performance of Signal Onset Method in Estimation of Arrival Time Delays.

| Method | Reference Signal | Bandpass Range | Mean of Abs. of Deviations ($ms$) | Mean of Deviations ($ms$) | $\sqrt{\text{Mean Square of TMHE}} (ms)$ |
| --- | --- | --- | --- | --- | --- |
| (1) Signal Onset | N/A | Tight | $10.8 \pm 67.9$ | $-1.00 \pm 71.6$ | $67.8 \pm 53.9$ |
| (2) Signal Onset | N/A | Loose | $13.6 \pm 81.5$ | $-2.00 \pm 86.2$ | $77.4 \pm 58.3$ |

\* TMHE := Truncated Mean Hydrophone Error

**Table 3.** Performance of Cross-Correlation Methods in Estimation of Arrival Time Delays.

| Method | Reference Signal | Bandpass Range<br>($kHz$) | Mean of Abs. of<br>Deviations ($ms$) | Mean of Devia-<br>tions ($ms$) | $\sqrt{}$ (Mean Square<br>of TMHE) ($ms$) |
| --- | --- | --- | --- | --- | --- |
| (3) Cross-Correlation | Hydro. 1 | Tight | $5.90 \pm 44.2$ | $-0.670 \pm 45.2$ | $48.0 \pm 64.0$ |
| (4) Cross-Correlation | Hydro. 1 | Loose | $12.9 \pm 22.8$ | $-1.70 \pm 27.0$ | $28.5 \pm 12.9$ |
| (5) Cross-Correlation | Source Signal | Tight | $6.5 \pm 80.6$ | $0.770 \pm 41.2$ | $38.7 \pm 48.0$ |
| (6) Cross-Correlation | Source Signal | Loose | $11.1 \pm 11.4$ | $2.6 \pm 16.3$ | $23.0 \pm 8.44$ |
| (7) Matlab “Find-Delay” | Hydro. 1 | Tight | $5.80 \pm 48.4$ | $-0.562 \pm 49.4$ | $58.3 \pm 79.4$ |
| (8) Matlab “Find-Delay” | Hydro. 1 | Loose | $12.8 \pm 22.8$ | $-1.8 \pm 27.0$ | $28.4 \pm 13.0$ |
| (9) Matlab “Find-Delay” | Source Signal | Tight | $6.50 \pm 40.0$ | $0.797 \pm 41.1$ | $38.7 \pm 48.0$ |
| (10) Matlab “Find-Delay” | Source Signal | Loose | $11.1 \pm 11.4$ | $2.70 \pm 16.3$ | $23.1 \pm 8.42$ |

\* TMHE := Truncated Mean Hydrophone Error

**Table 4.** Performance of GCC-PHAT Methods in Estimation of Arrival Time Delays.

| Method | Reference Signal | Bandpass Range<br>($kHz$) | Mean of Abs. of<br>Deviations ($ms$) | Mean of Devia-<br>tions ($ms$) | $\sqrt{}$ (Mean Square<br>of TMHE) ($ms$) |
| --- | --- | --- | --- | --- | --- |
| (11) GCC-PHAT | Hydro. 1 | Tight | $3.10 \pm 2.20$ | $-0.273 \pm 3.90$ | $6.48 \pm 2.29$ |
| (12) GCC-PHAT | Hydro. 1 | Loose | $3.10 \pm 2.20$ | $-0.273 \pm 3.90$ | $6.48 \pm 2.29$ |
| (13) GCC-PHAT | Source Signal | Tight | $239 \pm 354$ | $-143 \pm 422$ | $577 \pm 289$ |
| (14) GCC-PHAT | Source Signal | Loose | $227 \pm 344$ | $143 \pm 405$ | $568 \pm 304$ |

\* TMHE := Truncated Mean Hydrophone Error

**Table 5.** Performance of Spectrographic Cross-Correlation Methods in Estimation of Time.

| Method | Reference Signal | Bandpass Range<br>($kHz$) | Mean of Abs. of<br>Deviations ($ms$) | Mean of Devia-<br>tions ($ms$) | $\sqrt{}$ (Mean Square<br>of TMHE) ($ms$) |
| --- | --- | --- | --- | --- | --- |
| (15) 32-Sample Spec. Cross-Correlation | Hydro. 1 | Tight | $4.90 \pm 28.7$ | $-0.648 \pm 29.4$ | $17.2 \pm 18.6$ |
| (16) 32-Sample Spec. Cross-Correlation | Hydro. 1 | Loose | $5.90 \pm 6.10$ | $-0.939 \pm 8.8$ | $13.2 \pm 5.86$ |
| (17) 32-Sample Spec. Cross-Correlation | Source Signal | Tight | $10.7 \pm 65.5$ | $-1.10 \pm 68.4$ | $64.0 \pm 64.8$ |
| (18) 32-Sample Spec. Cross-Correlation | Source Signal | Loose | $22.5 \pm 45.3$ | $8.70 \pm 51.6$ | $54.8 \pm 37.4$ |
| (19) 64-Sample Spec. Cross-Correlation | Hydro. 1 | Tight | $5.10 \pm 29.2$ | $-0.383 \pm 30.0$ | $17.7 \pm 18.8$ |
| (20) 64-Sample Spec. Cross-Correlation | Hydro. 1 | Loose | $5.80 \pm 6.80$ | $-1.00 \pm 9.40$ | $13.5 \pm 6.78$ |
| (21) 64-Sample Spec. Cross-Correlation | Source Signal | Tight | $12.5 \pm 59.9$ | $-1.50 \pm 63.3$ | $70.7 \pm 72.8$ |
| (22) 64-Sample Spec. Cross-Correlation | Source Signal | Loose | $29.7 \pm 51.5$ | $11.0 \pm 60.7$ | $73.5 \pm 52.9$ |

\* TMHE := Truncated Mean Hydrophone Error
\* TMSE := Truncated Mean Sample Error

**Fig 6.**
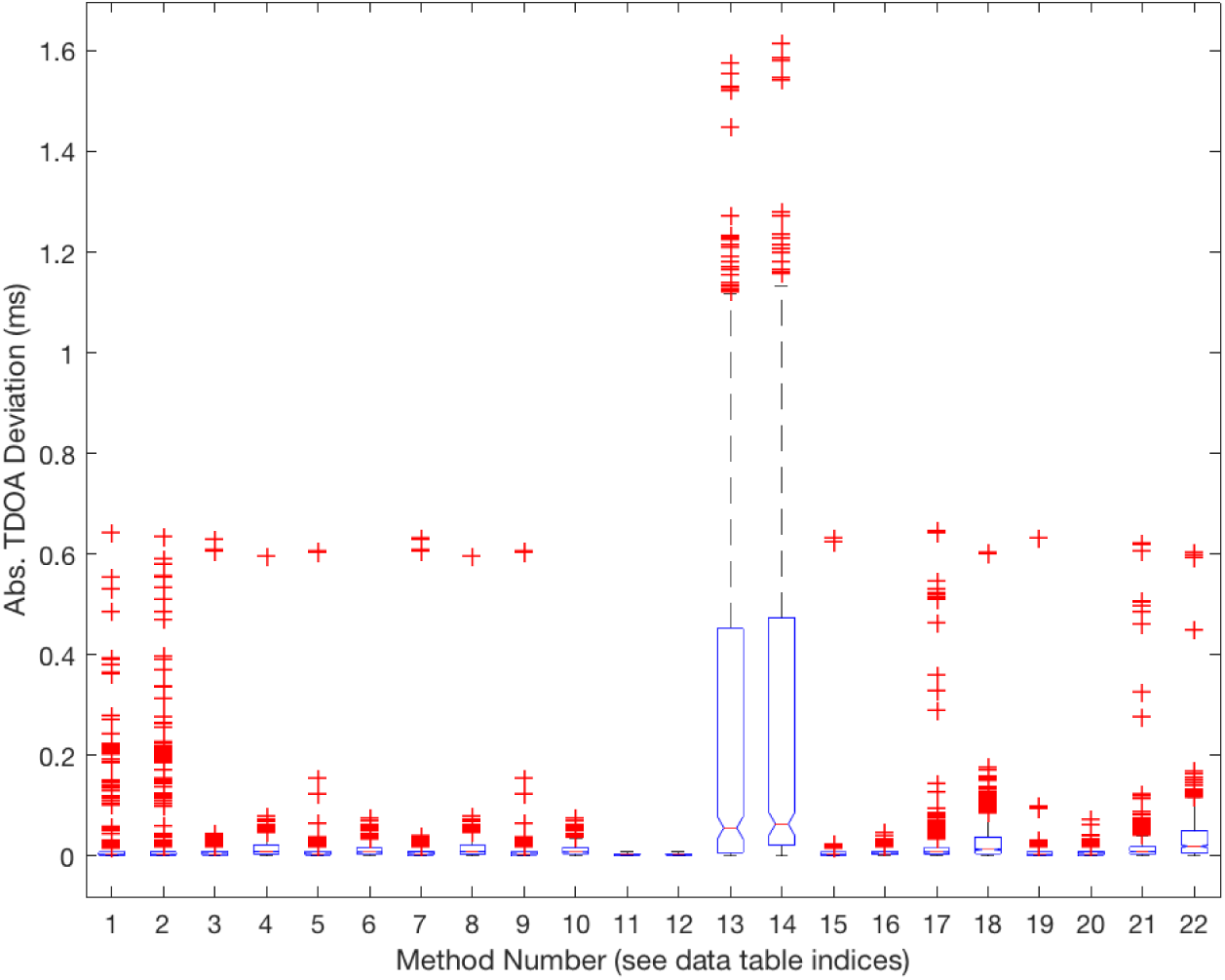
Standard box plot of absolute value TDOA deviations for all TDOA extraction methods.

**Fig 7.**
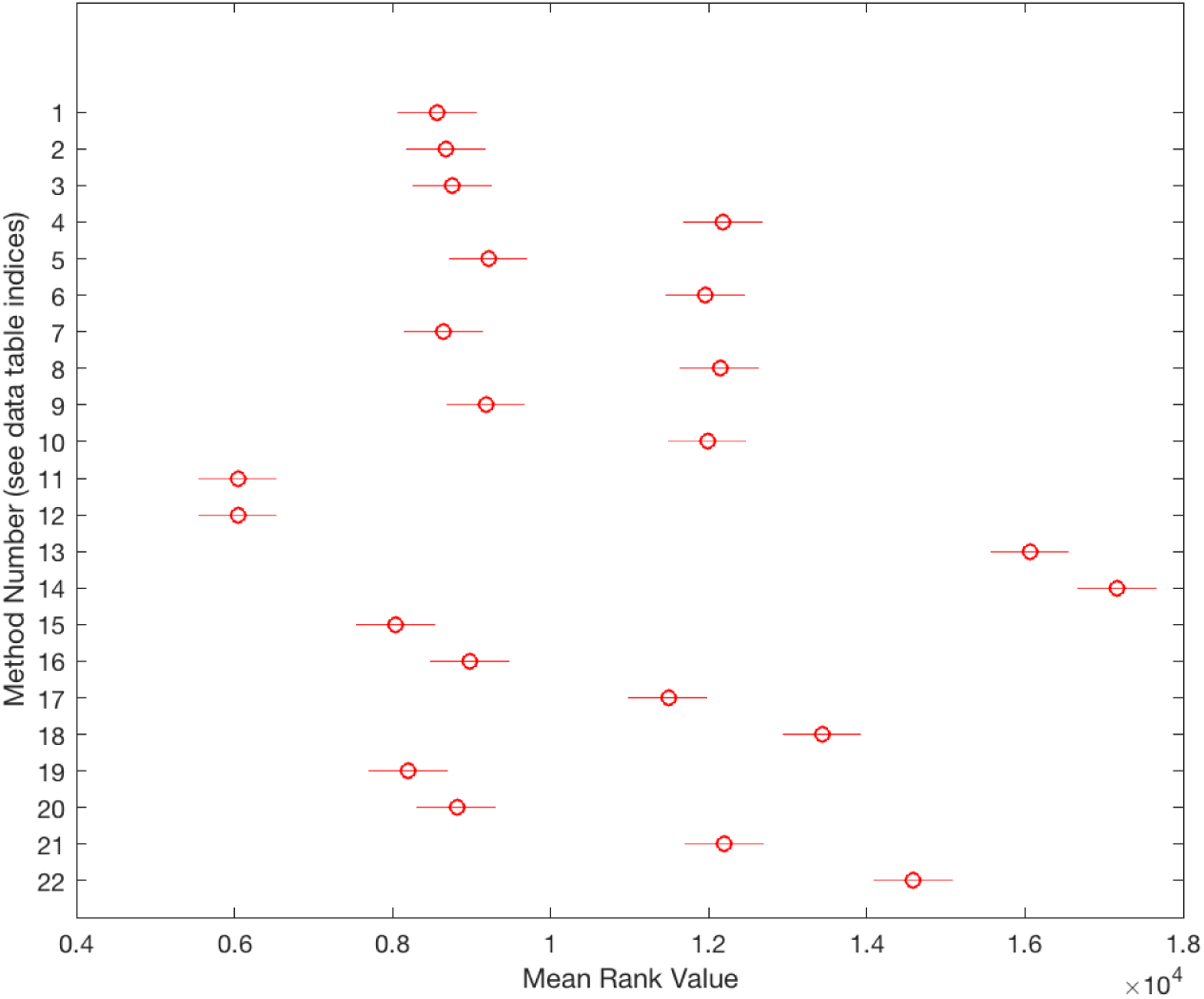
Mean rank intervals for all TDOA extraction methods.

### Whistle-like Sound De-noising

For each of the 14 unique play locations, we attempted standard and Weiner de-convolution of the artificial whistles using the corresponding IRF. A representative example of the results is shown in Fig 8. We determined that the increase in noise (and failure of the intended de-noising) was visually apparent enough to obviate the need for quantification.

**Fig 8.**
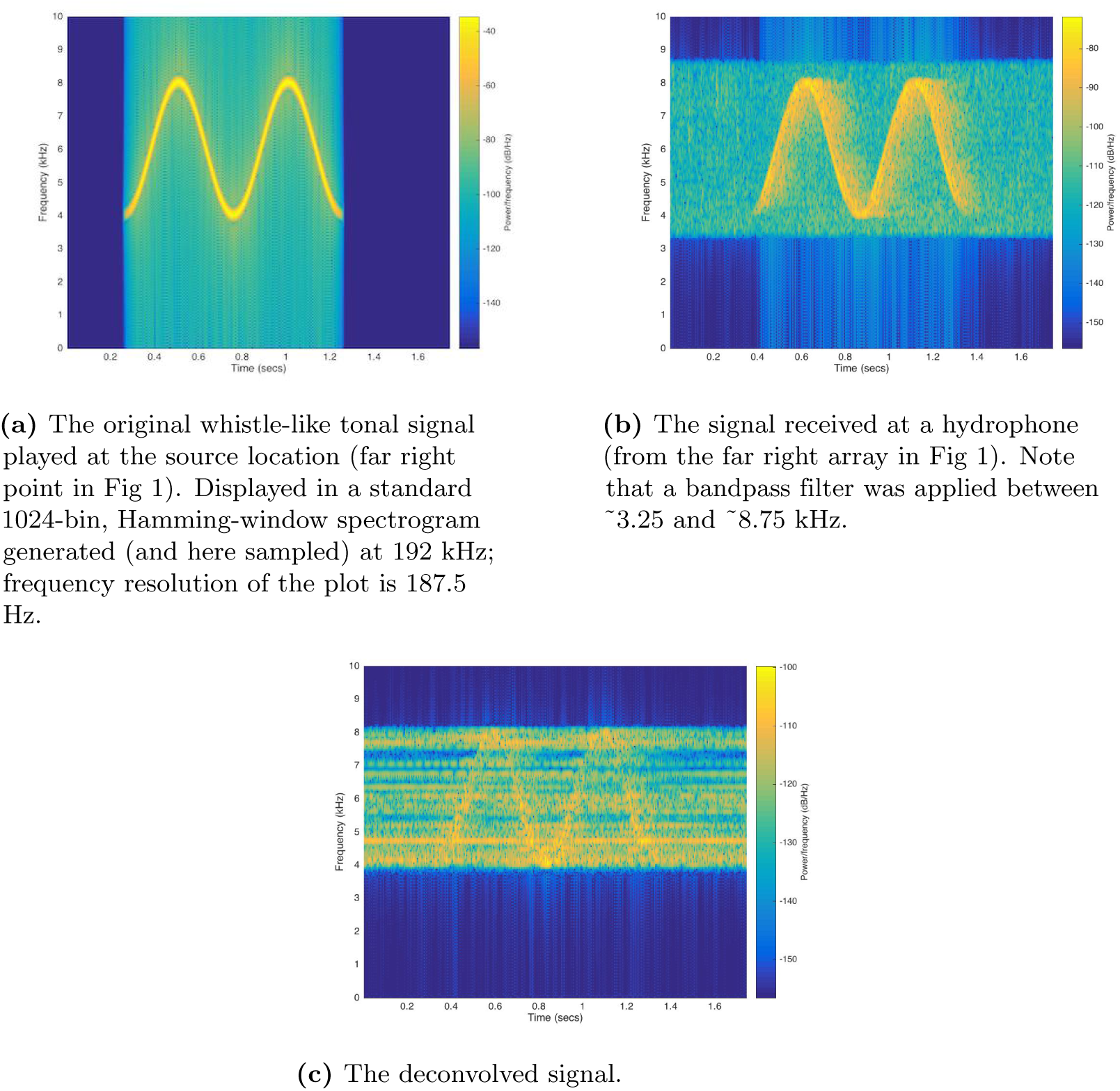
Wiener Deconvolution of a Whistle-like Signal with the IRF.

## Discussion

This paper represents an experimental performance evaluation of standard TDOA-based sound localization methods on both near-ideal impulses and whistle-like tones recorded by our custom audiovisual system, deployed in an unusually reverberant aquatic environment (a half-cylindrical dolphin pool). The first part of this paper is concerned with confirming that these methods, as implemented for our system, perform well for near-ideal impulses; since dolphin echolocation clicks have already been well localized in similar fashions [13–15], if at closer sensor-subject distances than we explore here, we would expect decent performance. The second part of this paper is concerned with evaluating these methods on tonal sounds modeled after *T. truncatus* whistles; they are theoretically expected to encounter difficulties [16] and, when experimentally evaluated individually in less rigorous circumstances, have performed modestly at best, not accomodating studies of acoustic exchanges [18–23]. In general, our results were what we expected: while near-ideal impulses were successfully localized, whistle-like tones were not.

First, we played “snow white” noise at 14 known locations in the pool, which allowed us to reconstruct IRF’s for those locations. By extracting time-differences-of-arrivals (TDOA’s) between pairs of sensors using a standard thresholding method (a common practice for localizing dolphin clicks, cited above), and subsequently feeding these TDOA’s into the standard Spherical Interpolation estimator to the system of equations describing the sound path geometry [9], we showed that the IRF’s could be localized with resolution comparable to that required for the practical purpose of separating two distinct dolphins: except close to the pool midline, error in accuracy was approximately half a mature bottlenose dolphin length, with error in precision longer (1-2 dolphin lengths) but confined primarily to a single axis. In general, these data seem to suggest that, were dolphins to vocalize signals resembling an impulse – their echolocation clicks might qualify – it would be possible to distinguish them at a separation of two or three body-lengths, depending on their relative orientation. Depending on dolphin number and clustering, this might certainly be adequate to achieve successful sound attribution for most vocalizations. It is important to note that cloud size was manually chosen to minimize the ratio of distance-to-point/area, and that there is room for a more rigorous quantitative optimization. Moreover, with a more substantial set of IRF’s it might be possible to develop a correction function that compensates for not only the localization clouds’ spread but their deviations from the expected source points.

As an aside, it is obvious that the cloud of localized points is always oriented towards the pool center, which is a result of the Spherical Interpolation method in combination with our sensor geometry that deserves further investigation. If it were possible to collapse the distribution with modifications in sensor geometry or the algorithm itself, the system’s capacity for sound localization might improve drastically.

We also played sounds constructed with parameters approximating those of *T. truncatus* whistles, detailed in our other work [27]. Even though they were not played from a speaker moving at average dolphin speed, the speed was not fixed, and so we argue these sounds constitute valid proxies for evaluating techniques for localizing dolphin whistles – even if we expect the techniques’ performance on these sounds to reflect the lower error bound of their performance on real whistles. We ultimately found that all of the techniques explored did not produce TDOA’s that provided any useful approximation of sound source localization using Spherical Interpolation. The TDOA errors exceeded approximately 3 ms, whereas the errors that generated the IRF spreads (Fig 4) were approximately 0.5 ms. As commented above, the unsuitability of these techniques for localizing tonal sounds in reverberant environments has been theoretically predicted and experimentally noted in less rigorous environments. Nevertheless, among the methods examined, from Fig 7 we note that maximization of the GCC-PHAT using a received signal as reference is the best method examined, regardless of whether the received signals were pre-processed with a tight or loose bandpass filter. We therefore assert that the GCC-PHAT is superior to the standard circular cross-correlation for the purposes of TDOA extraction in a reverberant environment.

Combining data from the two previous sections, we attempted to deconvolve the artificial whistles with corresponding IRF’s to obtain received signals cleansed of multipath effects, and thus false peaks in their cross-correlations. However, it immediately became clear that the signal-to-noise ratios of the IRF’s was not increased. This is most likely a result of the linear response not dominating the response function; in the future we would seek to obtain the full non-linear response. Also, although we chose our Lubbell LL916H speaker for its relatively flat frequency response, it is unclear whether it is adequately flat for the purposes of this calibration.

Overall, we have shown that, while our system installed at the Dolphin Discovery at National Aquarium is theoretically capable of localizing near-ideal, pulse-like sound in the pool using standard TDOA-based methodology, this methodology is not does not accomodate tonal sounds in this environment. Nevertheless, we note that GCC-PHAT significantly outperforms circular cross-correlation at the task of TDOA extraction, which might be relevant to beam-forming localization approaches. We expect our results to generalize to similar systems installed in similarly reverberant environments.

## Acknowledgments

We thank the National Aquarium for participating in this study, as well the National Science Foundation (Awards 1530544, 1607280), the Eric and Wendy Schmidt Fund for Strategic Innovation, and the Rockefeller University for funding. While regrettably we cannot name everyone, we also thank the approximately two dozen people at the National Aquarium, the Rockefeller University, and Hunter College for assisting with various aspects of the project.

## References

1. McBride AF, Herb DO. Behavior of the Captive Bottle-nose Dolphin, *Tursiops truncatus*. Journal of Comparative and Physiological Psychology. 1948;41(2):111–123.

2. Dreher JJ. Linguistic Considerations of Porpoise Sounds. The Journal of the Acoustical Society of America. 1961;33(12):1799–1800.

3. Caldwell MC, Caldwell DK. Individualized Whistle Contours in Bottlenosed Dolphins (*Tursiops truncatus*). Nature. 1965;207(1):434–435.

4. Caldwell MC, Caldwell DK. Vocalization of Naive Captive Dolphins in Small Groups. Science. 1968;159(3819):1121–1123.

5. Janik VM, Sayigh LS. Communication in bottlenose dolphins: 50 years of signature whistle research. Journal of Comparative Physiology A. 2013;199(6):479–489.

6. Caldwell MC, Caldwell DK, Tyack PL. Review of the signature-whistle-hypothesis for the Atlantic bottlenose dolphin. In: Leatherwood S, Reeves RR, editors. The Bottlenose Dolphin. San Diego; 1990. p. 199–234.

7. King SL, Sayigh LS, Wells RS, Fellner W, Janik VM. Vocal copying of individually distinctive signature whistles in bottlenose dolphins. Proceedings of the Royal Society B: Biological Sciences. 2013;280(1757):20130053–20130053.

8. Janik VM, King SL, Sayigh LS, Wells RS. Identifying signature whistles from recordings of groups of unrestrained bottlenose dolphins (*Tursiops truncatus*). Marine Mammal Science. 2013;29(1):109–122.

9. Li X, Deng ZD, Rauchenstein LT, Carlson TJ. Source-localization algorithms and applications using time of arrival and time difference of arrival measurements. Review of Scientific Instruments. 2016;87(4):041502–13.

10. Zimmer WMX. Passive Acoustic Monitoring of Cetaceans. Cambridge: Cambridge University Press; 2011.

11. Smith JO, Abel JS. Closed-Form Least-Squares Source Location Estimation from Range-Difference Measurements. IEEE Transactions on Acoustic, Speech, and Signal Processing. 1987;ASSP-35(12):1661–1669.

12. Smith JO, Abel JS. The Spherical Interpolation Method of Source Localization. IEEE Journal of Oceanic Engineering. 1987;OE-12(1):246–252.

13. Watkins WA, Schevill WE. Sound source location by arrival-times on a non-rigid three-dimensional hydrophone array. Deep Sea Research and Oceanographic Abstracts. 1972;19(10):691–706.

14. Watkins WA, Schevill WE. Listening to Hawaiian Spinner Porpoises, *Stenella* Cf. *Longirostris*, with a Three-Dimensional Hydrophone Array. Journal of Mammalogy. 1974;55(2):319–328.

15. Koblitz JC, Wahlberg M, Stilz P, Madsen PT, Beedholm K, Schnitzler HU. Asymmetry and dynamics of a narrow sonar beam in an echolocating harbor porpoise. The Journal of the Acoustical Society of America. 2012;131(3):2315–2324.

16. Spiesberger JL. Linking auto- and cross-correlation functions with correlation equations: Application to estimating the relative travel times and amplitudes of multipath. The Journal of the Acoustical Society of America. 1998;104(1):300–312.

17. Van Den Broeck B, Bertrand A, Karsmakers P, Vanrumste B, Van Hamme H, Moonen M. Time-domain generalized cross correlation phase transform sound source localization for small microphone arrays. In: Education and Research Conference EDERC, th European DSP; 2013.

18. Thomas RE, Fristrup KM, Tyack PL. Linking the sounds of dolphins to their locations and behavior using video and multichannel acoustic recordings. The Journal of the Acoustical Society of America. 2002;112(4):1692–1701.

19. Di Claudio ED, Parisi R. Robust ML wideband beamforming in reverberant fields. IEEE Transactions on Signal Processing. 2003;51(2):338–349.

20. Bell BM, Ewart TE. Separating Multipaths by Global Optimization of a Multidimensional Matched Filter. IEEE Transactions on Acoustic, Speech, and Signal Processing. 1986;ASSP-34(5):1029–1036.

21. Freitag LE, Tyack PL. Passive acoustic localization of the Atlantic bottlenose dolphin using whistles and echolocation clicks. The Journal of the Acoustical Society of America. 1993;93(4):2197–2205.

22. Janik VM, Thompson M. A Two-Dimensional Acoustic Localization System for Marine Mammals. Marine Mammal Science. 2000;16(2):437–447.

23. López-Rivas RM, Bazuá-Durán C. Who is whistling? Localizing and identifying phonating dolphins in captivity. Applied Acoustics. 2010;71(11):1057–1062.

24. Tyack PL. An optical telemetry device to identify which dolphin produces a sound. The Journal of the Acoustical Society of America. 1985;78(5):1892–1895.

25. Watwood SL, Owen ECG, Tyack PL, Wells RS. Signature whistle use by temporarily restrained and free-swimming bottlenose dolphins, Tursiops truncatus. Animal Behaviour. 2005;69(6):1373–1386.

26. Akamatsu T, Wang D, Wang K, Naito Y. A method for individual identification of echolocation signals in free-ranging finless porpoises carrying data loggers. The Journal of the Acoustical Society of America. 2000;108(3):1353–5.

27. Everest FA. Master Handbook of Acoustics. 4th ed. McGraw-Hill Professional; 2000.

28. Woodward SF, Reiss D, Magnasco MO. Machine Source Localization of *Tursiops truncatus* Whistle-like Sounds in a Reverberant Aquatic Environment. bioRxiv. 2019;(3):401–12.

29. Knapp CH, Carter C. The Generalized Correlation Method for Estimation of Time Delay. IEEE Transactions on Acoustic, Speech, and Signal Processing. 1976;24(4):320–327.

